# Synchronous processing of temporal information across the hippocampus, striatum, and orbitofrontal cortex

**DOI:** 10.1101/2024.11.10.622881

**Authors:** Akihiro Shimbo, Yukiko Sekine, Saori Kashiwagi, Shigeyoshi Fujisawa

## Abstract

Information processing for interval timing is supported by several brain regions, including the hippocampus, basal ganglia, and frontal cortical areas. However, little is known about the mechanism by which temporal information is processed cooperatively in the distributed brain network. Here, we investigated the neuronal processing of temporal information in the hippocampal CA1, dorsal striatum, and orbitofrontal cortex by simultaneously recording neuronal activity during a temporal bisection task. We found time cells representing elapsed time during the interval period across all three regions. Seeking a potential mechanism for the correlative representation of time, we found that theta oscillations were dominant in these areas and modulated the activity of these time cells. Moreover, the synchronization of the time cell pairs across the areas was also regulated by theta oscillations. Taken together, these results demonstrated the presence of synchronous activity of time cells across the three areas on a fine time scale, which was supported by theta oscillations. In addition, decoding analysis revealed that the activity of the time cells in these areas correlated with the rats’ decisions based on their internal time estimation, with the decoded time also showing correlations across the three regions. Thus, the cooperative activity of time-cell assemblies in the three regions reflected the recognition of elapsed time in the rats. In conclusion, these results demonstrate the pivotal role of neuronal synchronization of time cells in supporting temporal processing in the distributed brain network.

## INTRODUCTION

To adapt to complex environments, animals need to use a wide range of temporal information to appropriately adjust their behavior. In particular, temporal information on the scale of seconds is critical for anticipating the timing of an important event and preparing for timed behavior (Buhusi and Meck, 2005; Paton and Buonomano, 2018). Several brain regions, including the hippocampus, dorsal striatum (DS), and orbitofrontal cortex (OFC), have been proposed to be involved as neuronal circuit mechanisms for temporal processing on this scale of seconds (Buhusi and Meck, 2005; Petter et al., 2018). Recent studies have reported that hippocampal “time cells” fire at specific moments of a temporally organized event, encoding elapsed time information as sequential representation (Pastalkova et al., 2008; MacDonald et al., 2011; Kraus et al., 2013; Shimbo et al., 2021; Omer et al., 2023). Moreover, there is increasing evidence that time cells exist not only in the hippocampus, but also in the DS (Gouvea et al., 2015; Mello et al., 2015; Bakhurin et al., 2017; Toso et al., 2021) and the frontal cortical areas including the OFC (Bakhurin et al., 2017; Bolkan et al., 2017; Tiganj et al., 2017). Studies using lesion or pharmacological intervention techniques have also shown that these brain regions play essential roles in recognizing and processing information on interval timing. Previous studies suggest that lesioning the hippocampus or the disconnection of the fimbria-fornix alters animals’ anticipation of event occurrence earlier (Meck et al., 1984; Solomon et al., 1986; Tam and Bonardi, 2012) and hippocampal inhibition impaired rat’s discriminability between relatively similar length of durations (Jacobs et al., 2013). In addition, lesioning or pharmacological intervention of the DS disrupts interval timing (Meck, 2006; Macdonald et al., 2012) Accumulating evidence supports that the OFC contributes to the integration of reward size and delay length (Sosa et al., 2021).

These findings suggest that the temporal information is processed intrinsically by distributed neuronal circuits across various brain areas rather than by a single dedicated area (Paton and Buonomano, 2018). That is, temporal information is managed by correlational activity across related brain areas, rather than controlled by a central master clock in a single brain area. In this view, an important question is how distributed populations of neurons across these brain regions cope with shared temporal information. Previous studies have reported interregional correlations of neuronal activity across the brain areas in variety of cognitive functions (Jones and Wilson, 2005; Sjulson et al., 2018; Oberto et al., 2022). However, the mechanisms of synchronization of distributed temporal information across the CA1, DS, and OFC circuit for timed behavior remain to be elucidated.

To address this question, we investigated temporal processing in the dorsal hippocampal CA1, DS, and OFC by simultaneously recording neuronal activity and local field potentials (LFPs) during a temporal bisection task in which rats were required to measure and discriminate between long and short intervals in order to get a reward. We found time cells that encode elapsed time information during the interval period in the CA1, DS, and OFC. Bayesian decoding analysis revealed that rat estimates of elapsed time was able to be decoded similarly in these regions from the activity of time cells. Pursuing the mechanisms of correlative representations of elapsed time in these areas, synchronous structures of neuronal activity across these areas were examined. The results showed a dominance of the LFPs in the CA1, DS, and OFC with theta frequency of 4–10 Hz, and the LFPs were coherent across these areas. The spike timing of time cells in these areas was often significantly modulated by the theta oscillations. In addition, the synchronization of pairs of time cells across these areas was also modulated by the theta oscillations. These results suggest that neuronal synchronization of time cells in these three regions supported by theta oscillation may underlie the sharing of temporal information across the distributed brain network.

## RESULTS

### Representation of time in the CA1, DS, and OFC

Rats were trained on the temporal bisection task (Figure 1A and B; Material and Methods) (Shimbo et al., 2021). The tasks consisted of two types of trials: discrimination trials and test trials. In the discrimination trials, the rats were forced to run on a treadmill for long (10 s) or short (5 s) time intervals. The speed of the treadmill was randomly assigned as either 12 cm/s or 24 cm/s to dissociate elapsed time information from running distance. After the forced run, the rats had to choose the left or right arm in the Y-maze, which was associated with long- or short-time intervals, respectively. Water drops were provided as a reward at the end of the correct arm. Along with the discrimination trials of long- and short-time intervals, test trials in which the interval period was the geometric means of short and long ones (7.07 s) were randomly inserted. In the test trials, the rats also were required to choose the left or right arm of the Y-maze after forced running, although a reward and any sound stimuli were not contingent on the rats’ choice. There was no bias between the selection of the arms associated long- or short-time interval in the test trials (Figure 1C).

**Figure 1.**
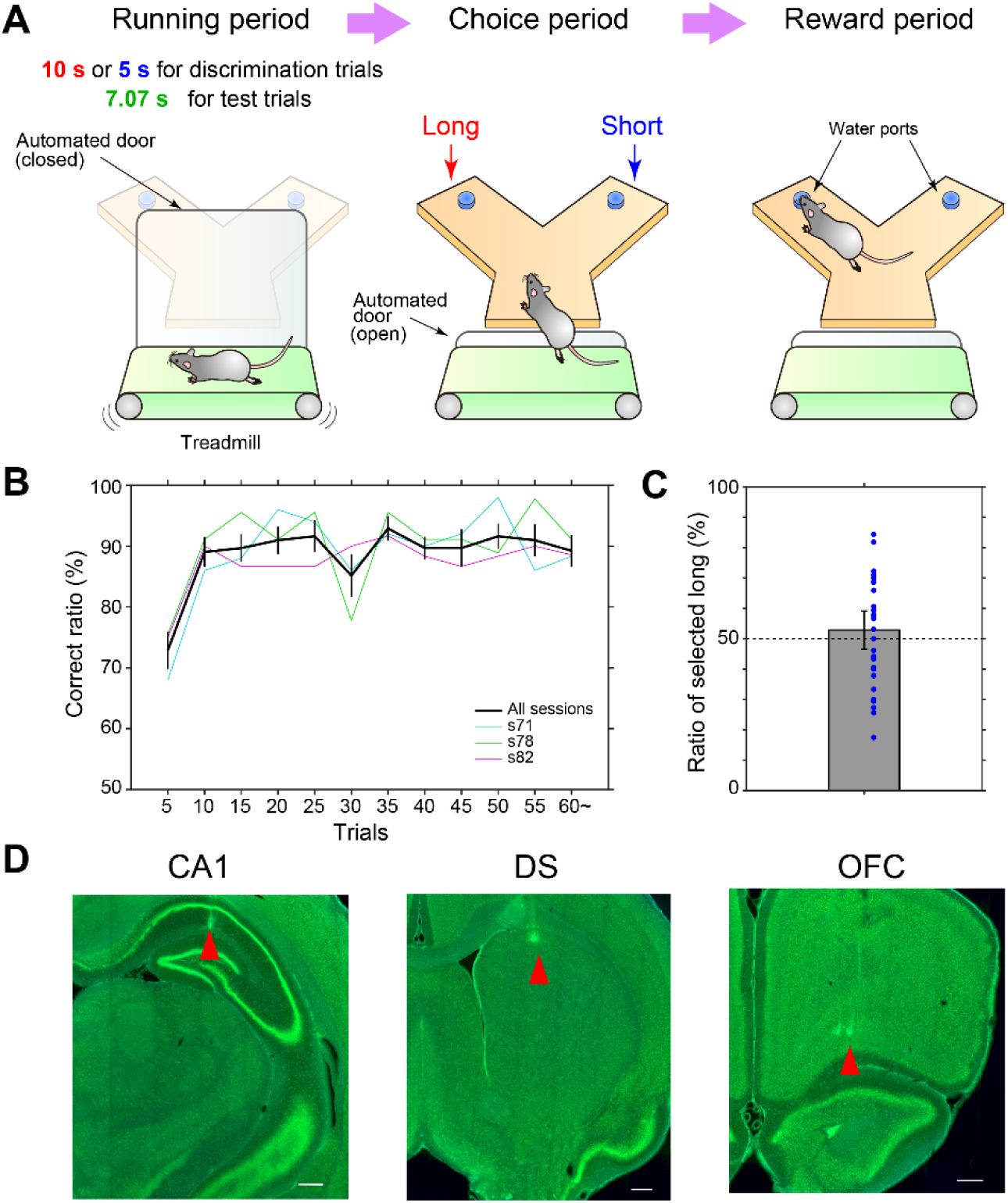
Recordings of neuronal activity from the CA1, DS, and OFC during the temporal bisection task. **(A)** Schematic of the temporal bisection task. Rats run on the treadmill for long (10 s) or short (5 s) time intervals. They then have to choose the left or right arm of the Y-maze, which are associated with long- or short-time intervals, respectively. A water drop is provided as a reward at the end of the correct arm. Test trials (7.07 s) are randomly inserted between the discrimination trials of short and long intervals. In the test trials, the rats are also required to choose the left or right arm of the Y-maze after forced running during the interval period, but no reward is provided in either arm. **(B)** Behavioral performance across sessions in the temporal bisection task. Performance of every five trials was averaged across sessions for all animals (black lines, mean ± S.E.M.; n = 31 sessions) or individual animals (colored lines, mean). **(C)** Average ratio of selected the arm associated with long-interval trials in the test trials in all sessions. The error bar indicates 95% confidence intervals (t-test). The ratio of each session is indicated by a blue dot. **(D)** Recording sites with silicon probes in the CA1, DS, and OFC of rat s71. Arrowheads, the traces of the probes on green fluorescent Nissl-stained sections. Scale bar: 0.5 mm.

Extracellular single units from the hippocampal CA1, DS, and OFC were simultaneously recorded during the task using high-density silicon probes (Figure 1D; Material and Methods). To examine the time-dependent firing activity of neurons in these areas during the interval period, peri-event time histograms (PETHs) of the discharges of single units during the discrimination trials were analyzed. A “time cell” was defined as a neuron that preferentially represented elapsed time information during the interval period using the following two criteria. First, units whose maximum firing rates were at least 2 Hz, and information rates (Skaggs et al., 1993) were at least 0.1 bit/spike during the long-time intervals were selected. Second, to identify neurons that preferentially coded temporal information (rather than distance or spatial information), the spiking activity of each neuron was fitted to generalized linear models with interval time, running distance, or spatial position as covariates (Kraus et al., 2013; Shimbo et al., 2021). Then, units whose spiking activity best fit the model with elapsed time compared to the other models with running distance or spatial position were selected (Material and Methods).

In total, 36.8% (300/815 units) of the CA1, 44.2% (254/575 units) of the DS, and 13.0% (172/1320 units) of the OFC neurons were classified as time cells according to the above two criteria (Figure 2A and B). The receptive fields of temporal information of the time cells in these areas were distributed over the interval period (Figure 2B). We then explored the similarities and differences in the coding characteristics of the time cells across these areas. First, the information rates of the time cells were analyzed. The information rates of time cells in the CA1 (0.51±0.43 bit/spike) and DS (0.58±0.43 bit/spike) were significantly higher than those in the OFC (0.21±0.14 bit/spike) (mean±SD; *F*_2,723_ = 53.1, *P* < 0.001, one-way analysis of variance (ANOVA) followed by Tukey-Kramer post-test; Figure 2C). Second, the extent of the receptive fields of the time cells in these areas was explored. The PETHs of the time cells were fitted with a Gaussian function, and the standard deviation (SD) of the Gaussian kernel was calculated (Material and Methods). The SDs of the time cells in the CA1 (1.39±0.88 s) and DS (1.44±0.85 s) were significantly smaller than those in the OFC (1.70±1.11 s) (mean±SD; *F*_2,723_ = 6.4, *P* < 0.01, one-way ANOVA followed by Tukey-Kramer post-test; Figure 2D). This result indicated that the receptive fields of time cells in the CA1 and DS were sharper than those in the OFC. Third, the peak firing rates of the time cells were analyzed, and the results showed that those of time cells in the CA1 (8.83±6.97 s) and DS (8.80±9.37 s) were significantly higher than those of time cells in the OFC (6.82±6.36 s) (mean±SD; *F*_2,723_ = 4.4, *P* < 0.05, one-way ANOVA followed by Tukey-Kramer post-test; Figure 2E).

**Figure 2.**
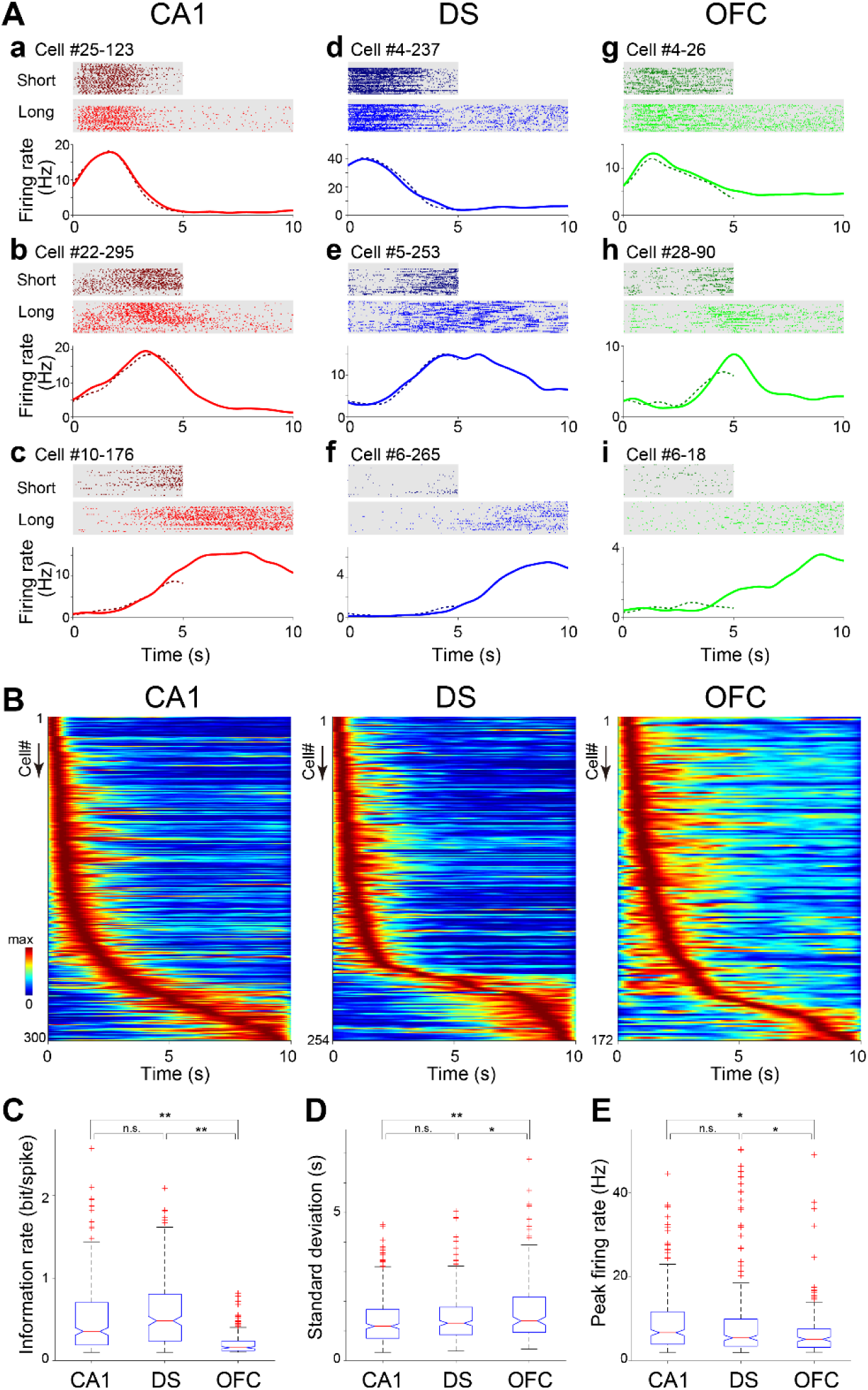
Representation of elapsed time in CA1, DS, and OFC neurons in the temporal bisection task. **(A, a-i)** Spiking activity of nine representative time cells in the neurons in the CA1 (a-c), DS (d-f), and OFC (g-i). *Top*, raster plots of time cells in long and short interval trials. *Bottom*, PETHs in long (solid line) and short (dotted line) interval trials. **(B)** Spiking activity of time cells in the CA1 (left), DS (middle), and OFC (right) (n = 298, 282, and 208 units from 3 rats, respectively). Each row represents the normalized PETHs of individual neurons in long-interval trials. The color scale shows the firing rate of each neuron (red represents the maximum rate of each neuron, and blue represents 0 Hz). Neurons are ordered according to the time of their peak firing rates. **(C)** Box plot of information rates of time cells in the CA1, DS, and OFC (n = 298, 282, and 208 units from 3 rats, respectively). **(D)** Box plots of standard deviation (SD) of Gaussian kernels fitted to PETHs of time cells in the CA1, DS, and OFC. **(E)** Box plots of maximum firing rates of PETHs of time cells in the CA1, DS, and OFC. \**P* < 0.05; \*\**P* < 0.01; ns, not significant. One-way ANOVA followed by Tukey-Kramer post-test.

In summary, there were time cells that encoded elapsed time information during the interval period in the CA1, DS, and OFC. Quantitatively, compared with the time cells in the OFC, those in the CA1 and DS had higher information rates, lower SD, and higher firing rates of receptive fields.

### Theta modulations of time cells in the CA1, DS, and OFC

To explore the mechanisms of synchronous representations of elapsed time information in the CA1, DS, and OFC, we next examined the interactions of LFPs across these areas. Wavelet analyses revealed that the LFPs in the CA1, DS, and OFC were dominated by a robust theta oscillation (4-10 Hz) during the interval period (Figure 3A and B). The wavelet coherence of the LFPs was also prominent in these areas (Figure 3C and D). These results suggested the presence of synchronous neuronal activity across the CA1, DS, and OFC during the interval period of the task. Therefore, we next investigated the mechanism by which spiking activity were modulated by theta oscillations in these areas (Figure 4).

**Figure 3.**
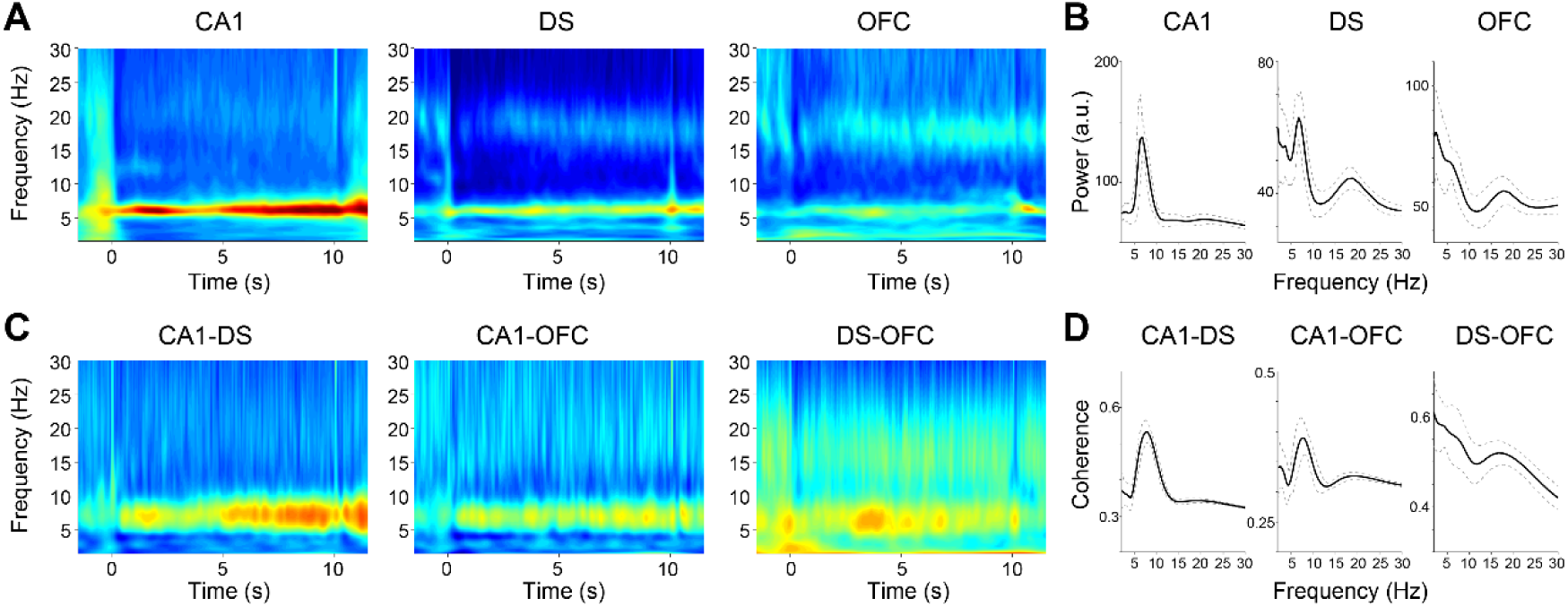
Theta modulations of LFPs in CA1, DS, and OFC during the temporal bisection task. **(A)** Time-resolved wavelet power spectrum of LFP in CA1, DS, and OFC during the interval period in a single rat (s82; mean of all sessions; n = 12 sessions). **(B)** Wavelet power spectrum of LFP in CA1, DS, and OFC during the interval period. Mean (solid line) ± SD (dashed line). n = 31 sessions from 3 rats. **(C)** Time-resolved wavelet coherence of LFPs between CA1 and DS, CA1 and OFC, and DS and OFC during the interval period in one rat (s82; mean of all sessions; n = 12 sessions). **(D)** Wavelet coherence of LFPs between CA1 and DS, CA1 and OFC, and DS and OFC during the interval period. Mean (solid line) ± SD (dashed line). n = 31 sessions from 3 rats.

**Figure 4.**
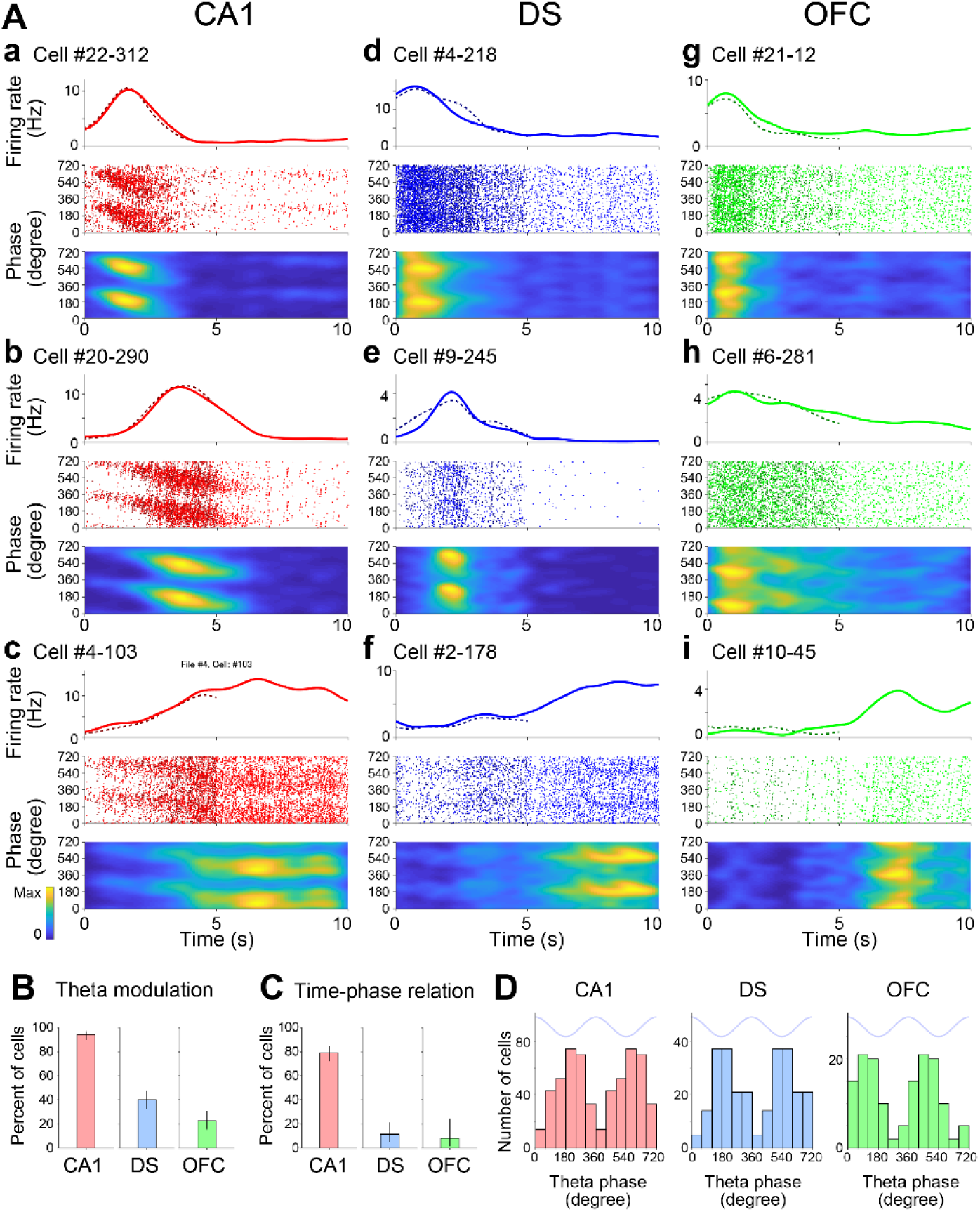
Entrainment of firing activity of time cells by theta oscillation. **(A,** a to i) Firing rates and theta phases of nine representative time cells in CA1, DS, and OFC. *Top*: PETHs of long (solid line) and short (dashed line) interval trials. *Middle*: Plots pf theta phase as a function of elapsed time. Each dot represents an action potential from the long (light color) and short (dark color) interval trials. *Bottom*: Histograms of theta phase plots as a function of elapsed time. **(B)** Ratios of time cells in CA1, DS, and OFC that are significantly phase modulated by hippocampal theta oscillations (*P* < 0.01, Rayleigh test). Clopper-Pearson confidence intervals for 99% are also shown. **(C)** Ratios of time cells in CA1, DS, and OFC that have significant time-phase correlations (*P* < 0.01, statistics of linear-circular correlation). Clopper-Pearson confidence intervals for 99% are also shown. **(D)** Distribution of preferred theta phases for significantly modulated (*P* < 0.01) time cells in CA1, DS, and OFC.

The spiking activity of 94.0% (99% confidence interval (CI): [89.6, 97.0]), 39.0% (99% CI: [31.2, 47.2]), and 21.5% (99% CI: [14.0, 30.6]) of CA1, DS, and OFC neurons, respectively, were significantly phase-modulated by the hippocampal theta oscillation (*P* < 0.01, Rayleigh test; Figure 4A and B). CA1 neurons are known to exhibit theta phase precession,(O’Keefe and Recce, 1993; Royer et al., 2012; Shimbo et al., 2021) so we examined whether there were phase-time relationships of individual neurons in these areas. The results showed that 80.9% (99% CI: [74.1, 86.5]) of the theta-modulated time cells in the CA1 showed phase precession in the interval period (Figure 4A and C; Material and Methods). Meanwhile, 8.1% (99% CI [2.7, 17.8]) of the theta-modulated neurons in the DS showed a significant phase-time relationship, which was slightly above chance level. In the OFC, 8.1% (99% CI [0.9, 26.6]) of the theta-modulated neurons showed a significant phase-time relationship, with CI overlapping the chance level. The preferred phases of significantly modulated neurons were similarly distributed for the CA1, DS, and OFC, although the phases of highest density were slightly shifted among the areas (Figure 4D). In summary, the majority of the CA1 time cells exhibited robust theta modulation with phase precession. In contrast, a large portion of the time cells in the DS and OFC were phase-locked to theta oscillation without phase precession.

### Theta modulations of synchronization of time cells within and across the areas

Given that sizable populations of the time cells in the CA1, DS, and OFC displayed robust theta modulation, we hypothesized that theta oscillation underlies the synchronization of the time cells across these areas. First, to evaluate interactions of time cells at fine time scales, the cross-correlograms (CCGs) of spike times of neuronal pairs were calculated within and across the CA1, DS, and OFC. This method allowed the assessment of the co-occurrence of spikes of the neuronal pairs at short latencies, indicating synchronous activity of the neuronal pairs. The results showed that some portions of the CCGs of time cell pairs within and across the CA1, DS, and OFC had sharp peaks at short latencies (∼100 ms) (Figure 5A). Jitter techniques were used to examine whether a peak of the CCG of a neuronal pair is significantly higher than that expected from the firing rates of the neurons (Fujisawa et al., 2008; Amarasingham et al., 2012). In this method, each spike in each neuron was randomly jittered in the original data set within ±100 ms, and a cross-correlogram with a surrogate dataset was computed. The process was repeated 1,000 times. Short-latency peaks in the original CCGs were determined to be statistically significant when they were higher than those constructed from the jittered data sets. The results showed that sizable portions of CCGs of neuronal pairs within and across these areas had significant peaks at short latencies (Figure 5B and C; 815 of 1806 pairs of CA1-CA1 time cells, 392 of 2120 of DS-DS, 96 of 680 of OFC-OFC, 362 of 2565 of CA1-DS time cells, 110 of 1658 of CA1-OFC, and 130 of 2229 of DS-OFC; *P* < 0.01, jitter method).

**Figure 5.**
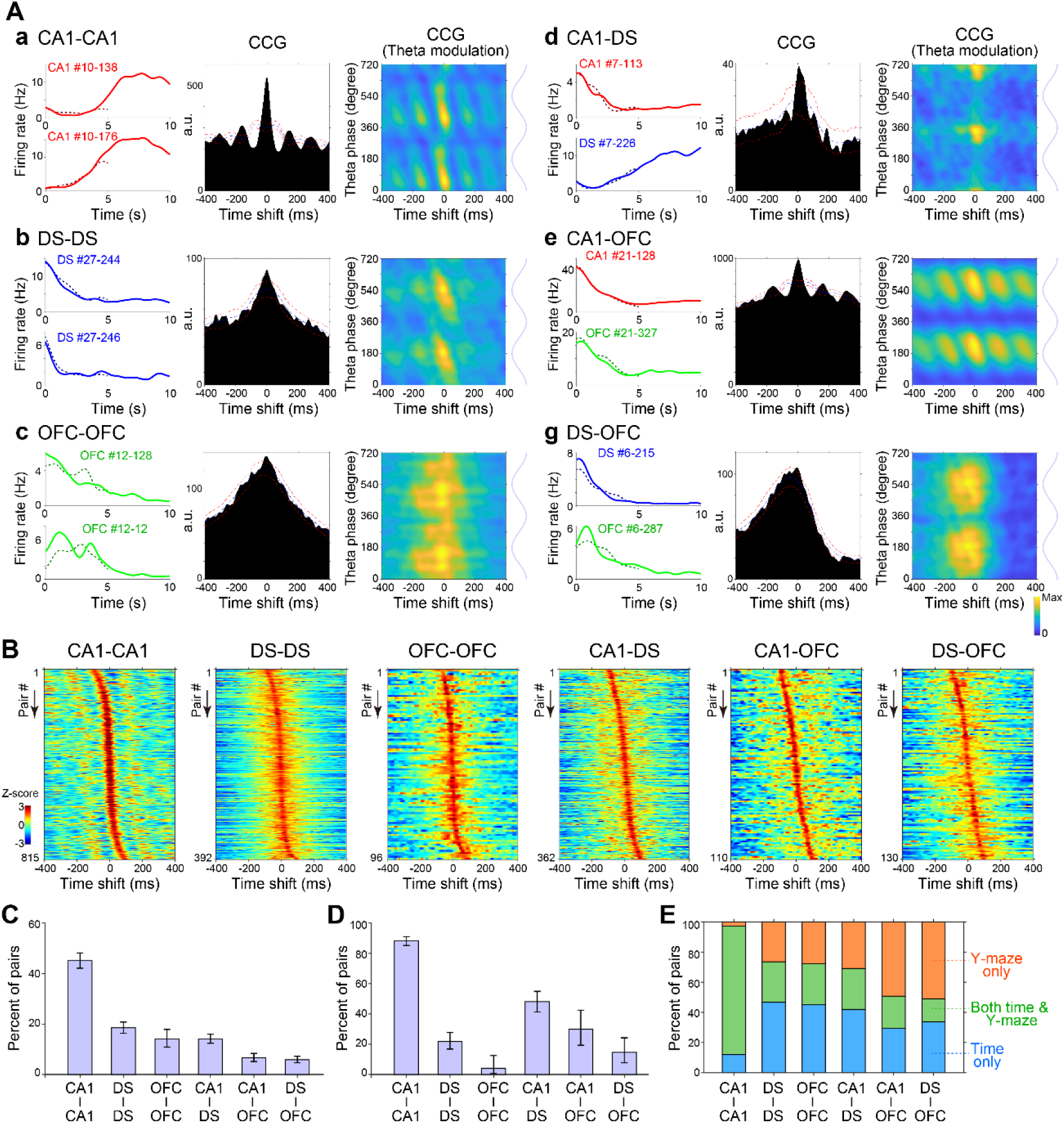
Synchronous activity of time cells within and across the CA1, DS, and OFC. **(A, a–g)** Synchronous activity of six representative time-cell pairs within and across the CA1, DS, and OFC. *Left*: PETHs of long (solid line) and short (dashed line) interval trials of the pairs of time cells. *Middle*: Cross-correlograms (CCGs) of the pairs of the time cells (jitter methods). *Right*: CCGs of neuronal pairs as a function of theta phase. **(B)** CCGs of time-cell pairs showing significant peaks within and across the CA1, DS, and OFC. The CCGs are z-scored in the plots. **(C)** Ratios of time-cell pairs with significant peaks within and across the CA1, DS, and OFC (jitter methods). Clopper-Pearson confidence intervals for 99% are also shown. **(D)** Ratios of time-cell pairs that are significantly modulated by theta oscillations within and across the CA1, DS, and OFC (Rayleigh test). Clopper-Pearson confidence intervals for 99% are also shown. **(E)** Ratios of time-cell pairs that have significant peaks within and across the CA1, DS, and OFC (jitter methods) during running on the treadmill only (time only; bule), running in the Y-maze only (Y-maze only, orange), and both treadmill and Y-maze (both time & Y-maze; green).

Second, we investigated whether the synchronization of time cells was modulated by theta oscillation by examining how cross-correlograms of time-cell pairs were phase-modulated by theta oscillations. CCGs of example neuronal pairs as a function of the theta phase were plotted (Figure 5A), and the plots showed that the CCG peaks were clearly modulated by the theta phase. To statistically evaluate phase-modulations of spike co-occurrence in neuronal pairs, we examined whether synchronous spike events (within ±50 ms) were phase-modulated by theta oscillations, using the Rayleigh test for non-uniformity of circular data. Sizable portions of CCGs of neuronal pairs within and across these areas had significant peaks at short latencies (720 of 815 pairs of CA1-CA1 time cells, 86 of 392 of DS-DS, 4 of 96 of OFC-OFC, 174 of 362 of CA1-DS, 33 of 110 of CA1-OFC, and 19 of 130 of DS-OFC; *P* < 0.01; Figure 5D). Taken together, these results demonstrated the presence of synchronous activity of time cells within and across the CA1, DS, and OFC on a fine timescale of ∼100 ms, which was supported by phase modulations of hippocampal theta oscillations.

In addition, we investigated whether the synchronous activity of time cells was organized only during treadmill running or also during running in the Y-maze (Figure 1A). To this end, we applied CCG analysis to the time cell pairs also during the Y-maze period. The results showed that some populations of pairs had significant peaks in their CCGs only in the treadmils (time only), while other population had significant peaks in both treadmils and Y-maze (both time & Y-maze) or only in the Y-maze (Y-maze only) (102, 713, and 25 pairs of CA1-CA1 time cells, 249, 143, and 141 of DS-DS, 249, 143, and 141 of OFC-OFC, 249, 143, and 141 of CA1-DS, 249, 143, and 141 of CA1-OFC, and 249, 143, and 141 of DS-OFC were time only, both time & Y-maze, and Y-maze only, respectively; Figure 5E). The result suggests that the synchronization of time cells within and across the CA1, DS, and OFC was organized in a behavioral phase-dependent manner during the temporal bisection task.

### Activity of time cells in the CA1, DS, and OFC represent rat decisions

Next, we investigated how the activity of time cells in the CA1, DS, and OFC were related to the internal recognition of elapsed time in the rats. For this purpose, the activity of the time cells was analyzed in the test trials (Figure 1A). In the test trial, rats were forced to run the interval time (7.07 s), which was the geometric means of long (10 s) and short (5 s). After the running interval, the rats needed to select either the arms for the long or short one. Here, we hypothesized that the rat selection in the test trials would reflect their decision about short- or long-time intervals depending on their internal estimation of elapsed time. Thus, we examined whether the activity of the time cells actually manifested the choices of the rats in the test trials by taking advantage of a Bayesian decoding technique. Previous studies demonstrated that with this technique, positional or temporal information of rats can be decoded from the spiking activity of their place cells or time cells, respectively (Brown et al., 1998; Zhang et al., 1998; Davidson et al., 2009; Pfeiffer and Foster, 2013; Feng et al., 2015; Terada et al., 2017; Danjo et al., 2018; Shimbo et al., 2021).

The Bayesian decoding technique was used to estimate the elapsed time information encoded by the time-cell assemblies in the CA1, DS, and OFC (Material and Methods). First, the spike trains of time cells that were recorded simultaneously from these areas were divided into 20-ms time bins in each trial. Second, the probability densities of elapsed time information from the spike counts of the time cells in each time bin were estimated using the Bayesian decoding method. The PETHs of the time cells during the discrimination trials were used for the decoding templates. Figure 6A shows an example of elapsed time information decoded with neurons of the CA1, DS, and OFC at each time bin in a single discrimination trial. As shown in the figure, decoded time information from spiking activity was well correlated with the actual elapsed time in the trial. The differences in the decoded time information between the test trials in which the rats selected the left (“selected-long”) or the right arm (“selected-short”) were then analyzed (Figure 6B).

**Figure 6.**
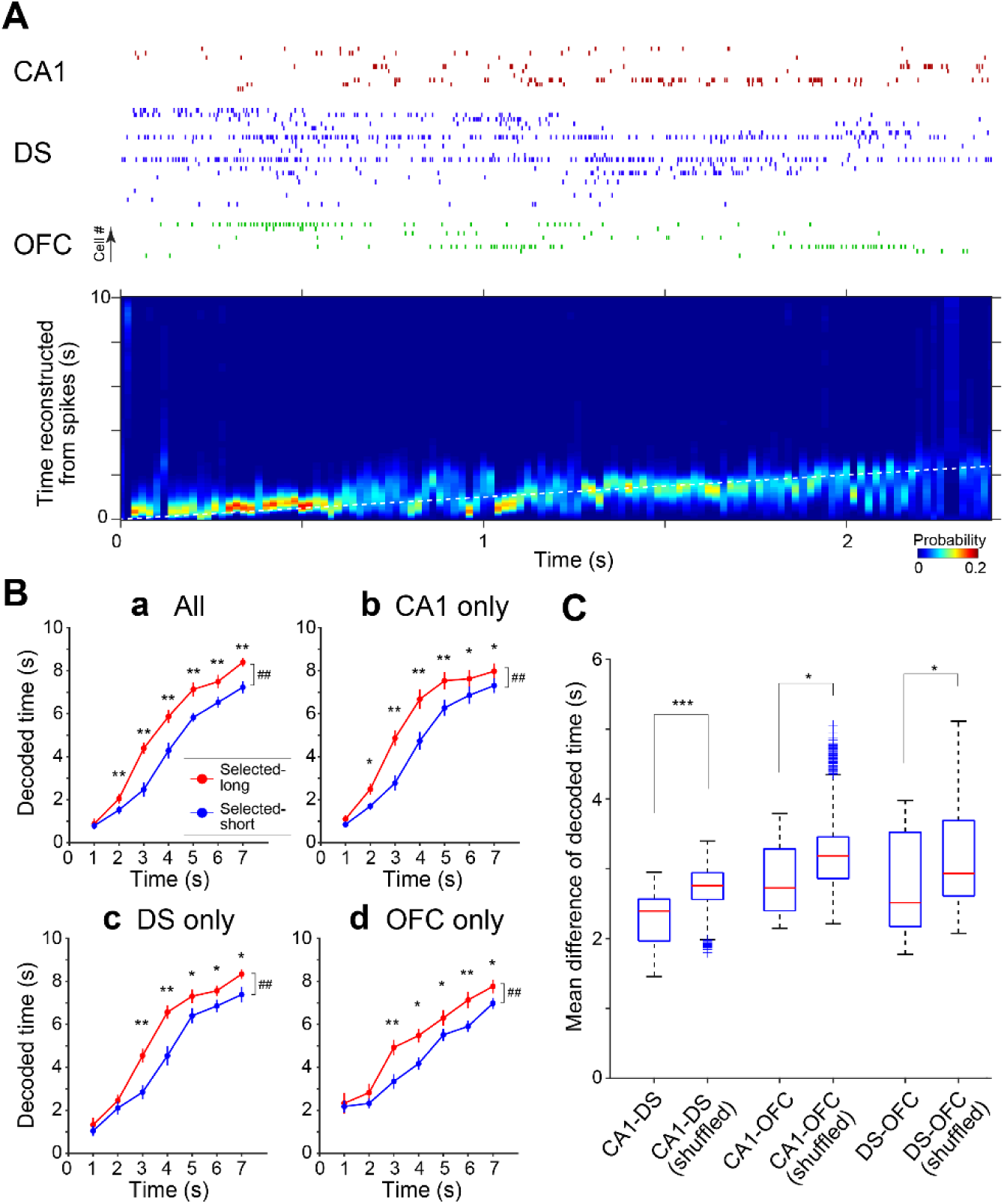
Activity of time cells in CA1, DS, and OFC reflects the decisions of rats. **(A)** Representative data of Bayesian decoding analysis. *Top*, simultaneously recorded spiking activity of time cells (11, 22, and 8 units from CA1, DS, and OFC, respectively) in a single normal trial of the temporal bisection task. *Bottom*, the probability density of temporal information is computed from the simultaneously recorded activity of time cells in the trial. **(B, a-d)** Decoded time in the selected-long and -short test trials per 1 s interval duration using spiking activity in CA1, DS, and OFC (a), only CA1 (b), only DS (c), and only OFC (d). Mean ± S.E.M. ##*P* < 0.01 for effects of trial type. \**P* < 0.05, \*\**P* < 0.01 for simple effects of trial type at each period. Two-way ANOVA with Bonferroni post-test. **(C)** Mean differences in decoded time across CA1, DS, and OFC in each trial. \**P* < 0.05, \*\*\**P* < 0.001, t-test compared to trial-shuffled data.

The elapsed time information of the selected-long and selected-short trials was calculated separately from the spiking activity of all of the CA1, DS, and OFC during every 1-second duration in the interval period in each session. If activity of time cells were related to the estimation of time in rats, decoded time from selected-long trial was larger than that from selected-short trial. This was because rat selection of arm in the test trials depended on the similarity between estimated time in the given test trial and long- or short-time interval. There were significant differences in decoded temporal information between the selected-long and selected-short trials (Figure 6Ba; n = 14 sessions from three rats; 2-way ANOVA followed by Bonferroni post-test; effect of trial type, *F*_1,13_ = 57.77, *P* < 0.001; interaction of trial type and period, *F*_6,78_ = 6.26, *P* < 0.001).

We applied the same decoding analysis using spike trains from each of CA1, DS, or OFC separately in the same trials. The results showed significant differences in decoded temporal information between the selected-long and selected-short trials in CA1, DS, or OFC (n = 14 sessions from three rats; 2-way ANOVA followed by Bonferroni post-test; CA1: effect of trial type, *F*_1,13_ = 47.49, *P* < 0.001; interaction of trial type and period, *F*_6,78_ = 8.03, *P* < 0.001 (Figure 6Bb); DS: effect of trial type, *F*_1,13_ = 26.24, *P* < 0.001; interaction of trial type and period, *F*_6,78_ = 5.27, *P* < 0.001 (Figure 6Bc); OFC: effect of trial type, *F*_1,13_ = 21.32, *P* < 0.001; interaction of trial type and period, *F*_6,78_ = 2.06, *P* = 0.07 (Figure 6Bd)).

We also examined whether decoded time information was correlated across these brain areas within each trial. In this analysis, the elapsed time information was decoded separately from the spiking activity of all of the CA1, DS, and OFC during every 1-second duration in the interval period in each trial, and the differences of the encoded times across these brain areas were estimated. As for statistical comparison, we also calculated the differences of the decoded times across these brain areas with shuffling the trials. The result showed that the time differences across these areas were significantly smaller than those of trial-shuffled data (Figure 6C).

These results demonstrate that the activity of time-cell assemblies in the CA1, DS, and OFC cooperatively predicted rat decisions during the test trials, indicating that the CA1, DS, and OFC commonly reflected the recognition of elapsed time in the rats.

## DISCUSSION

This study investigated the mechanism by which temporal information is recognized and processed in the distributed network of the CA1, DS, and OFC. Neuronal activity in the CA1, DS, and OFC was simultaneously recorded during the temporal bisection task, and the results showed that large fractions of neurons in these areas represent elapsed time information during the interval period of the task. To explore the mechanisms of correlative representations of elapsed time in these areas, synchronous structures of neuronal activity across the areas were also investigated. Wavelet analysis of the LFPs revealed that theta-frequency oscillation was dominant and coherent across the CA1, DS, and OFC. The activity of time cells in these areas was also modulated by theta oscillations. Furthermore, the synchronization of time cell pairs across the CA1, DS, and OFC was also modulated by theta oscillations. We applied the Bayesian decoding analysis to the activity of these time cells during the test trials and found that the decoded temporal information from the neurons of all these areas was correlated to the choices of the rats. This indicates that the activity of the time cells of these areas similarly represented rat decisions based on their internal estimation of the time. These results demonstrated that neuronal synchronization of time cells across the CA1, DS, and OFC supported by theta oscillation existed during the behavioral task requiring time measurement.

### Temporal processing in the CA1, DS, and OFC

Information processing of interval timing in the range of seconds to minutes is implemented in several brain regions, including the hippocampus, basal ganglia, and prefrontal cortex (Buhusi and Meck, 2005; Petter et al., 2018). The hippocampus is well known to be involved in encoding spatiotemporal information, such as event sequence, to support episodic memory (Burgess et al., 2002; Fortin et al., 2002; Terada et al., 2017; Shahbaba et al., 2022). Recent studies have shown that time cells in the CA1 reliably represent temporal information during task behavior (Pastalkova et al., 2008; MacDonald et al., 2011; Kraus et al., 2013; Salz et al., 2016; Shimbo et al., 2021; Omer et al., 2023). Importantly, time cells in the CA1 have similar coding properties to those of place cells, such as theta phase precession and scalability of representation (Pastalkova et al., 2008; Shimbo et al., 2021). This suggests that time cells may organize cognitive maps for temporal information as place cells form map representations for spatial navigation (Eichenbaum, 2014). Lesion studies also support the involvement of the hippocampus in time estimation functions for interval timing (Meck et al., 1984; Tam and Bonardi, 2012; Yin and Meck, 2014).

The basal ganglia have also been studied as an important site for the recognition and representation of time (Buhusi and Meck, 2005; Paton and Buonomano, 2018). Behavioral and physiological studies have provided evidence for the involvement of the basal ganglia in sensory timing and time perception (Harrington et al., 1998; Meck et al., 2008). Similar to the hippocampus, neurons in the striatum display representations of elapsed time information and organize sequential activity (Gouvea et al., 2015; Mello et al., 2015; Toso et al., 2021). A recent study by Monterio et al showed that temperature manipulations in the striatum affected the timing of activity in time cells and influenced the animals’ decision in temporal discrimination tasks (Monteiro et al., 2023), suggesting the role of temporal patterns of sequential activity in timing behavior.

The prefrontal cortex, including the orbitofrontal cortex, is thought to be involved in temporal integration functions.(Fuster, 2001; Xu et al., 2014; Parker et al., 2015; Tiganj et al., 2017; Wang et al., 2018) Prefrontal neurons also exhibit representations of elapsed time information (Tiganj et al., 2017), and temperature manipulations of the prefrontal cortex also affect animal’s estimates of elapsed time (Xu et al., 2014). Even when limited to the OFC alone, previous studies have reported the involvement of the OFC in various types of cognitive functions that require temporal information processing, such as the integration of temporal information into value-based decisions (Sosa et al., 2021). Physiological studies have also demonstrated the presence of time cells in the OFC (Bakhurin et al., 2017).

These collective findings indicate that several brain regions, including the hippocampus, basal ganglia, and frontal cortex, are involved in time perception for interval timing, suggesting a distributed nature of temporal information processing (Buhusi and Meck, 2005; Paton and Buonomano, 2018). The current study mainly focused on the encoding of temporal information in the CA1, DS, and OFC. Consistent with the previous studies, we found neurons encoding elapsed time information during the interval period in the CA1, DS, and OFC. While individual time cells in these areas played specific roles in processing of elapsed time information, populations of the time cells fired relatively uniformly throughout the entire interval period. Decoding analysis further revealed that activity in time cells within the CA1, DS, and OFC accurately represented the rats’ internal estimates of elapsed time. It is worth noting that debate persists regarding the mechanisms underlying internal time perception, highlighting the importance of considering the relationship between motor behavior and temporal perception (Robbe, 2023). Nonetheless, our findings suggest that neuronal encoding of temporal information is distributed across these brain regions.

### Neuronal synchronization in the CA1, DS, and OFC supported by theta oscillation

From the viewpoint of distributed encoding, the fundamental question is the neuronal circuit mechanisms of correlative encoding of elapsed time information among the involved brain areas. Previous research has reported interregional correlations of neuronal activity across brain areas in various types of cognitive functions (Engel and Singer, 2001; Buzsáki, 2006). In particular, it has been proposed that hippocampal theta oscillation plays a pivotal role in brain network communication (Battaglia et al., 2011). Theta oscillations in the hippocampus occur during active states of the brain, such as running or exploration. Theta oscillations are also dominant in a wide range of brain areas, including the neocortex and subcortical structures, and contribute to the dynamic formation of cell assemblies across the regions (Siapas et al., 2005; Sirota et al., 2008; Benchenane et al., 2010; Colgin, 2011). Thus, it has been proposed that theta oscillations support online information processing in cognitive behaviors.

Regarding the interaction between the hippocampus and striatum, gamma oscillations and neuronal spiking activity in the striatum are often phase-modulated by hippocampal theta oscillations in a behavior-dependent manner (Berke et al., 2004; DeCoteau et al., 2007; Tort et al., 2008) In the frontal cortical areas including the OFC, neuronal activity is also dynamically modulated by theta oscillations in various cognitive behaviors (van Wingerden et al., 2010; Benchenane et al., 2011; van Wingerden et al., 2014; Jarovi et al., 2018; Guillem and Ahmed, 2020; Knudsen and Wallis, 2020, 2022). Importantly, the involvement of theta oscillations in temporal perception has also been proposed in previous studies. A lesion of the medial septum, which is an important site of theta oscillation generators (Buzsaki, 2002) induces a shift of initiation of timed behavior to an earlier direction (Meck et al., 1987). In addition, the sequential activity of CA1 time cells is strongly disrupted after infusion of the GABA_A_ agonist muscimol into the medial septum (Wang et al., 2015). Taken together, these results support that theta oscillations would be a possible substrate for synchronization of the distributed network to support time perception and recognition.

The current study investigated whether and how hippocampal theta oscillations would underlie neuronal synchronization for sharing temporal information across the distributed brain areas. Wavelet analysis of LFPs revealed the dominance of the theta frequency band in CA1, DS, and OFC, which was coherent across these areas. In addition, the spike timing of time cells in these areas was significantly phase-modulated by theta oscillations. We also examined neuronal interactions within and across the areas by analyzing cross-correlations of spike timing of the neuronal pairs. This analysis was performed because interregional synchronous neuronal activity is thought to underpin dynamic communication across related brain areas to support various cognitive functions (Benchenane et al., 2010). The results showed that sizable portions of the time cell pairs within and across the CA1, DS, and OFC were significantly synchronized on fine time scales of ∼100 ms (Figure 6), indicating the existence of functional connectivity of time cells across the distributed areas. Importantly, these correlations of spike times of time cell pairs were often phase-modulated by theta oscillations. Collectively, these results suggest contributions of theta oscillations to interregional neuronal communication for temporal perception and recognition.

In conclusion, the results of this study demonstrated that the neurons in the CA1, DS, and OFC commonly encode temporal information that represented the rat’s internal recognition of time. The activity of these time cells in these areas was modulated by theta oscillations, which may promote flexible interregional neuronal interaction on fine time scales. Such neuronal synchronization of time cells across the areas underlies the possible mechanisms of temporal information sharing in the distributed network.

## MATERIALS AND METHODS

### Key resources table

| Key Resources Table |  |  |  |  |
| --- | --- | --- | --- | --- |
| Reagent type (species) or resource | Designation | Source or reference | Identifiers | Additional information |
| strain, strain background (Mus musculus) | Long-Evans rat | Japan SLC | Iar:Long-Evans | male |
| software, algorithm | Matlab | Mathworks | N/A | https://www.mathworks.com/ |
| software,<br>algorithm | Kilosort2 | (Pachitariu<br>et al., 2024) | N/A | https://github.com/MouseLand/Kilosort |
| software,<br>algorithm | DeepLabCut | (Mathis et<br>al., 2018) | N/A | https://github.com/DeepLabCut/DeepLabCut |
| Other | Silicon probe: 64-site, 6-shank | NeuroNexus | Buzsaki64sp | For<br>electrophysiology<br>recordings |
| Other | 256-channel<br>Multiplexed<br>Biosignal<br>Amplifier | Amplipex,<br>Ltd. | KJE-1001 | For<br>electrophysiology<br>recordings |

### Subjects

Three male Long-Evans rats (aged 8–10 weeks at the start of training) were used in this study. All experimental protocols were approved by the RIKEN Institutional Animal Care and Use Committee.

### General behavioral methods

Six-week-old male Long-Evans rats were purchased from Japan SLC, Inc. (Shizuoka, Japan). The rats were housed in a temperature-controlled room (20–22℃) with a 12-h light-dark cycle (lights were turned on at 8:00 and off at 20:00). When the rats weighed more than 250 g, a water deprivation protocol was started in which the amount of water intake was restricted to 10 ml outside of the task, but the food was available *ad libitum*. Training on the temporal bisection task (8-10 weeks old at the start of training) started 2 days later from the beginning of the water deprivation schedule.

The apparatus for the behavioral experiments were described in our previous report (Shimbo et al., 2021). Briefly, the behavioral experiments were conducted on the apparatus consisting of a treadmill (W15 × D40 cm) and Y-maze (20 cm length for the central arm and 24 cm length for the side arms), which were partitioned by a transparent acrylic automated door (Figure 1A). The treadmill was surrounded by transparent acrylic walls (50 cm in height) except for the door side (right side to the running direction). Two water ports for rewards were placed at the ends of the arms of the Y maze, and one water port was placed on the front walls of the treadmill. These water ports had tube connections with micro-pumps (Type 7615; Christian Bürkert GmbH & Co. KG, Ingelfingen, Germany) that delivered 0.2% saccharin water (M5N1991; Nacalai tesque, Kyoto, Japan) as a reward. Two LED lights (OXL/CLH/80 Series; Oxley, Singapore) were positioned at the end of the arm for light stimuli. For the system control, four infrared photoelectric sensors (PZ-G51P; Keyence, Osaka, Japan) were set for detection of the rat positions in the apparatus; one sensor was placed on the treadmill, one was set at the entrance of the Y-maze, and two were located at a 12 cm distance from the end of the arm. Behavioral experiments were automatically controlled using the custom-written LabVIEW (National Instruments, Austin, TX) programs running on a Windows PC.

### Temporal bisection task

Three rats (rat-id: s71, s78, and s82) were trained for the temporal bisection task (Figure 1A) (Shimbo et al., 2021). The tasks consisted of two types of trials, discrimination trials and test trials. In the discrimination trials, the rats were required to discriminate between long (10 s) and short (5 s) intervals on the treadmill. In test trials, the rats were presented with the geometric means of the short and long intervals (7.07 s). Discrimination and test trials randomly appeared in a single session.

The experimental procedures of a single discrimination trial were as follows. When the rats entered the treadmill, the door was closed, and a small water reward (15 µl) was provided at the water port on the wall of the treadmill with a sound stimulus (8 kHz sine wave, 1 s). After the sound presentation, the treadmill started to move, and the rats were forced to run for long (10 s) or short (5 s) intervals. The speed of the treadmill was randomly assigned in each trial. After running, the small reward at the water port on the wall was provided with the sound stimulus (8 kHz sine wave, 1 s) again. Concurrently, the door opened, and the LED lights on both arms of the Y-maze were turned on. The rats were required to select either the left or right arm in the Y-maze, which were associated with long- or short-time intervals, respectively. If the rats selected the correct side, the LED lights were turned off, the sound stimulus (4 kHz sine wave, 1 s) was presented, and a water reward (20 µl) was provided at the water port of the correct arm. If the rats selected the incorrect side, the LED lights were turned off, and a different sound stimulus (11 kHz square wave, 1 s) was presented; however, the rats did not receive a water reward. After an incorrect response, the correction trials, which consisted of the same sets of intervals and speeds as the incorrect trials, were performed in the subsequent trial.

The experimental procedures of the test trial were similar to those of the discrimination trial. In the test trials, however, the interval period for running on the treadmill was always the geometric means of the short and long intervals (7.07 s). The rats also had to select the left or right arm of the Y-maze after forced running during the interval period, but rewards and sound stimuli were not provided in either of the arms. The reason for using geometric means of the short and long intervals is that previous studies reported that bisection points are located around the geometric means of intervals (Church and Deluty, 1977; Lejeune and Wearden, 2006).

The schedule for training and recording sessions was as follows. After the rats had habituated to the apparatus, training sessions started. In the training sessions, only the discrimination trials were performed. The speed of the treadmill was set to either 12, 18, or 24 cm/s. One session consisted of 60 trials in which the correction trials were not included or a duration of 60 min.

After the rats showed more than 80% correct response rates for three successive sessions (not including the correction trials), the silicon probes were implanted surgically. Following the postoperative recovery period (4–7 days), the water deprivation procedure and training session were started again. When the rats reached stable behavioral performances again (more than 80% correct response rate for two consecutive sessions), a recording session started. A recording session consisted of discrimination and test trials described above. The speed of the treadmill was set to either 12 cm/s or 24 cm/s. The session consisted of 80 trials. Four sets of intervals and speeds in the discrimination trials (2 intervals × 2 speeds) were presented pseudo-randomly in 60 trials. Two sets of intervals and speeds in the test trials (1 interval × 2 speeds) were inserted in 33% probability. Neural activity was analyzed when rats showed a correct response rate of more than 80% during both the long and short trials (excluding the correction trials and test trials).

### Surgery and recording

After rats showed good performance in the training session (rats became around 14–32 weeks old), we implanted silicon probes in three rats (rat-id: s71, s78, and s82) for the chronic recording of neuronal activity during the tasks. General surgical procedures for chronic recordings have been described previously (Terada et al., 2017). In this study, Buzsáki64sp silicon probes (NeuroNexus, Ann Arbor, MI) were used for recording. The probes consisted of 6 shanks (200-µm shank separation). Each shank had ten recording sites (160 µm^2^ each site; ∼1 MΩ impedance), staggered to provide a two-dimensional arrangement (20 µm vertical separation). The rats were implanted with three silicon probes in the left hemisphere of the OFC (the targeted coordinate was anterior-posterior (AP) = +4.2 mm, medial-lateral (ML) = −1.6 mm, and dorsal-ventral (DV) = − 4.0 mm), the DS (AP = +0.2 mm, ML = −2.4 mm, and DV = −3.0 mm), and the CA1 (AP = − 5.0 mm, ML = −3.7 mm, and DV = −2.1 mm). The silicon probes were placed approximately 0.5 mm above the targeted coordinates at the time of surgery and inserted gradually daily to the desired depth positions with the attached micromanipulator.

During the recording sessions, the wide-band neurophysiological signals were recorded continuously at 20 kHz on a 256-channel Amplipex system (KJE-1001; Amplipex Ltd, Szeged, Hungary) (Berenyi et al., 2014). Kilosort2 was performed for spike sorting, and then the experimenter manually adjusted the clusters (Pachitariu et al., 2024). For LFP analysis, signals downsampled to 1.25 kHz were used. Rat behavior was monitored with a USB camera with a CMOS sensor (acA720-520; Basler AG, Ahrensburg, Germany) set on the ceiling. The capturing times of video frames were simultaneously recorded with electrophysiological data to synchronize the physiological and behavioral data in the *post hoc* analyses. The positional information of the head of the rat was subtracted from the video frames using the DeepLabCut (Mathis et al., 2018) software.

After all of the recording sessions were finished, a small current (5 µA for 10 to 12 s) was passed through the recording electrodes of the probe 2 days prior to sacrificing the animals to identify the recording sites. The rats were deeply anesthetized and perfused through the heart first with 0.9% saline solution followed by 4% paraformaldehyde solution. The brains were sectioned by a slicer (PRO10; Dosaka EM, Kyoto, Japan) at 100 µm in the coronal plane. Sections were stained with fluorescent Nissl, mounted on slides, and cover-slipped. The slides were scanned on a NanoZoomer Digital Pathology HT (Hamamatsu Photonics) with ×20 resolution. The tracks of the silicon probe shanks were reconstructed from multiple sections (Figure 1B).

### Data analysis

All analyses after spike sorting were performed using custom-written programs in MATLAB (MathWorks, Natick, MA). We used the Circular Statistics Toolbox (Berens, 2009) for circular analyses and statistics.

### Selection of time cells

A time cell was defined as a unit that preferentially represented the information of elapsed time during the interval period, using the following selection criteria. First, to identify units that represented temporal information during the interval period, we selected the units whose maximum firing rates were at least 2 Hz and information rates (Skaggs et al., 1993) were at least 0.1 bit/spike during the interval period in the task. Second, to determine units that preferentially encoded temporal information rather than distance or spatial information, we fitted the spiking activity of each neuron to a generalized linear model (GLM), with elapsed time, running distance, or spatial position as the covariates (Kraus et al., 2013; Shimbo et al., 2021). We selected units whose spiking activity was best fitted to the model that used elapsed time as the covariate as compared with the other models that used running distance and spatial position as the covariates. The GLM analysis is detailed below.

### GLM analysis

The GLM analysis was based on our previous work (Shimbo et al., 2021). To estimate the effects of elapsed time, running distance, and spatial position on unit firing during the interval period, we applied a GLM to the spiking activity of each neuron (MacDonald et al., 2011; Kraus et al., 2013). We modeled the unit discharges during the interval period as an inhomogeneous

Poisson process with the firing rate as a function of covariates (elapsed time, distance, or position) that modulates spiking activity (MacDonald et al., 2011; Lepage et al., 2012; Kraus et al., 2013). To evaluate the effect of each covariate, the following three models were compared. The “time” model is a fifth-order polynomial of elapsed time *τ*(*t*) described as

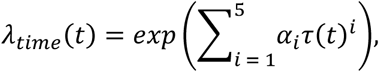

where *α*_*i*_ are parameters to fit. The “distance” model was a fifth-order polynomial of running distance on the treadmill (*d*(*t*)) described as

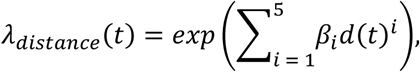

where *β*_*i*_ are parameters to fit. The “position” model is a Gaussian-shaped place field that is described as

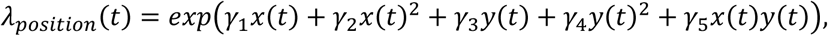

where *γ*_*i*_ are parameters to fit. The spiking activity of each neuron was fitted to each model. The interval period was divided into 20 ms bins, and spike counts of the unit in each bin were estimated from the real data. We used the *glmfit* function of MATLAB for this analysis to fit the spiking data to each model and compared the deviances of the fit across the models. The best model was selected by comparing the deviances of the fit.

### Bayesian decoding method for reconstructing temporal information from unit activity

The Bayesian decoding analysis was based on our previous work (Shimbo et al., 2021). A memoryless Bayesian decoding algorithm (Brown et al., 1998; Zhang et al., 1998) was used to estimate elapsed time information during the interval period from the spike trains of time cells in the CA1, DS, and OFC (Figure 3). Based on the Bayes’ theory, the posterior probability of elapsed time from a trial onset (‘time’) given spike trains from single neurons (‘spikes’) was described as

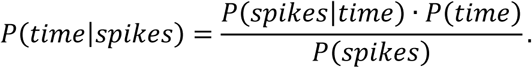

The prior probability is described as follows, under the assumption that spiking events follow a Poisson distribution and that firing rates of neurons are independent of each other:

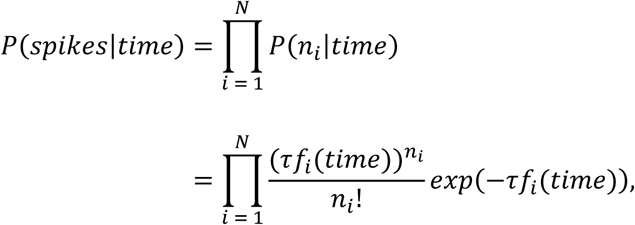

where τ is the time window (20 ms) of sampling spike trains, *f*_*i*_(*time*) is the PETH of *i-*th neuron, *n_i_* is the number of spikes of *i-*th neuron in the time window, and *N* is the total number of neurons. Combining these equations, the posterior probability of time is described as

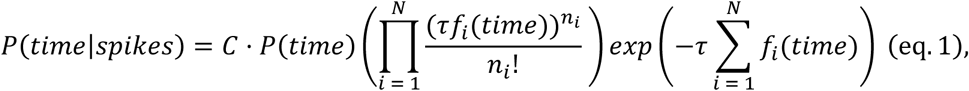

where *C* is a normalization factor that depends on τ and the numbers of spikes of each neuron (Zhang et al., 1998).

Taking advantage of the Bayesian decoding method described above, we analyzed differences in neuronal activity between the test trials in which the rats selected the arm that was correct in the long-interval trails (“selected-long”) and the arm that was correct in the short-interval trials (“selected-short”) (Figure 3B). The posterior probability *P*(*time*|*spikes*) (eq. 1) was estimated separately in the selected-long and selected-short test trials. *f*_*i*_(*time*) was used as the PETH of *i-*th unit estimated in the discrimination trials and *n_i_* as the number of spikes of *i-*th unit in the test trials. To assess differences in the decoded time information between selected-long and selected-short test trials, we first calculated the probability densities of elapsed time of the selected-long and selected-short trials from the spiking activity during every 1-second duration in the interval period in each session. Then, the average of the peak times of the two types of trials in each 1-second duration was computed. The two-way analysis of variance followed by Bonferroni post-test using R software (Team, 2013) was used to statistically compare the data between selected-long and selected-short trials. In this analysis, the sessions in which four or more-time cells were simultaneously recorded in each area (CA1, DS, and OFC) were selected (n = 14 sessions).

### Theta phase modulation and phase precession

For the theta phase extraction, the LFPs in the CA1 pyramidal cell layers, which were determined by the amplitudes of ripples and the polarities of sharp waves (Mizuseki et al., 2011), were filtered with a Butterworth filter with a pass-band range of 4–10 Hz. Instantaneous theta phases were estimated using the Hilbert transformation of the filtered signals. To assess whether the neuronal activity was phase-modulated with theta oscillations, the Rayleigh test was used to assess the circular uniformity of spike events of a neuron on theta phases (Figure 5B) (Sirota et al., 2008; Berens, 2009). For analyzing theta phase precession, the relationship between the elapsed time and theta phase of spiking activity in each unit during the interval period was analyzed. The presence of theta phase precession was determined according to a significant linear-circular correlation (*P* < 0.01) between time and theta phases during the interval period.(Berens, 2009)

### Cross-correlation analysis of neuronal pairs

The cross-correlograms (CCGs) of the cell pairs within and across the CA1, DS, and OFC were analyzed to assess the synchronous activity of neuronal pairs. Jitter techniques were used to examine whether a peak of the CCG of a neuronal pair was significantly higher than that expected from the firing rates of the neurons (Fujisawa et al., 2008; Amarasingham et al., 2012). We randomly jittered each spike in each neuron in the original data set within ±100 ms and computed a cross-correlogram with a surrogate dataset. The process was repeated 1,000 times, and then, short-latency peaks in the original cross-correlogram were determined to be statistically significant when they were atypical with respect to those constructed from the jittered data sets (*P* < 0.01).To statistically evaluate phase-modulations of spike co-occurrence in neuronal pairs, we examined whether synchronous spike events (within ±50 ms) were phase-modulated by theta oscillations, using the Rayleigh test for non-uniformity of circular data (*P* < 0.01).

## AUTHOR CONTRIBUTIONS

A.S. and S.F. designed the experiments. A.S., Y.S. and S.K. performed experiments. A.S. and S.F. analyzed the data. A.S. and S.F. wrote the paper.

## ACKNOWLEDGMENTS

The authors thank RIKEN Advanced Manufacturing Support Team for supporting the fabrication of the custom-made experimental apparatus. We also thank the Support Unit for Bio-Material Analysis, Research Resources Division, RIKEN Center for Brain Science, for technical support. This study was supported by the JSPS KAKENHI Grant (18H02711 and 18H05525 for S.F.; 20K16480 for A.S.).

## Notes

### Competing Interest Statement

The authors have declared no competing interest.

